# Stage-specific and tomato-spotted wilt virus infection-induced changes in the salivary gland transcriptome of western flower thrips

**DOI:** 10.64898/2026.08.26.745926

**Authors:** Joshua B. Benoit, Sulley Ben-Mahmoud, Swapna Priya Rajarapu, Christopher J. Holmes, Samuel T. Bailey, Diane Ullman, Dorith Rotenberg

## Abstract

Western flower thrips (WFTs) are critical vectors of tomato spotted wilt virus (TSWV), transmitting it via a circulative-propagative cycle. The insect-virus relationship is unusual in that only larvae can acquire the virus for transmission to plants to occur. During the larval stage, the virus circulates and replicates within many organs, reaching the salivary glands before the insect pupates, and remaining in infected organs when the insect becomes an adult. The virus continues to replicate in the salivary glands of adult insects, after which it is inoculated into plants via saliva during feeding. Understanding the interactions between TSWV and the WFT salivary glands is critical to furthering investigations of TSWV inoculation and efforts to block the spread of this devastating plant virus. Here, we document transcriptomic changes associated with TSWV infection of the salivary glands of adults (males and females) and second instar larvae. Gene sets enriched in adult male, female, and larval genes revealed a core set of genes associated with WFT salivary glands, as well as genes that differed between sexes and between adults and larvae. The transcriptome response to TSWV infection was higher in larvae (second instar in this study) than in adults, with nearly a 10x increase in differentially expressed genes. We hypothesize this occurred because larvae efficiently acquire the virus and the virus first enters the SGs at the L2 stage, whereas adult SGs are infected only if acquisition occurred in the larval stage. Thus, assessment of larvae detects responses to the early stages of infection, while assessment of adults detects responses to the later stages of infection. Similarly, functional changes in larval salivary glands were more diverse, with significant transcriptome differences associated with growth and development in this tissue during infection. Lastly, a comparative analysis of changes in a published SG proteome revealed a correlation between transcript and protein levels during infection, but little overlap between significant TSWV-responsive transcripts and proteins. These studies provide critical insight into the molecular changes associated with the first breach of the SGs in larvae by TSWV, revealing a markedly different transcriptomic response compared to that in adults.

## Introduction

The Western flower thrips (WFT), *Frankliniella occidentalis* (Pergande) (order Thysanoptera), is a globally distributed agricultural insect pest that causes significant economic losses through feeding, oviposition, and the transmission of several plant-infecting viruses (Reitz et al. 2020). WFTs are polyphagous, known to feed on hundreds of species of plants, colonizing, feeding and laying eggs on foliage, stems and flowers of a broad range of hosts (Childers & Achor 1995). Adults are omnivorous, also feeding on non-plant hosts, including fungi and spider mite eggs (Stafford-Banks et al. 2014).

Host plants can be cultivated in field and/or greenhouse environments and include diverse vegetables, fruit trees, and ornamentals, including important crops such as tomatoes, peanuts, lettuce, peppers, roses, and orchids (Hollingsworth & Armstrong 2005; Reitz et al. 2020; Demirozer et al. 2012). Using their piercing-and-sucking mouthparts, WFTs puncture into plant tissues, emptying cells and ingesting the sap (Hunter & Ullman 1994). This feeding behavior, along with the habit of emptying cell contents, causes characteristic scars and silvery patches on the foliage and fruits of infested plants (Kindt et al. 2003). Under heavy infestation, the destruction from feeding alone is enough to result in diminished yield, distorted and unmarketable fruit, or unappealing ornamental plants/flowers. Thrips transmission of orthotospoviruses, such as tomato spotted wilt virus, undoubtedly causes the greatest damage to crops (Batuman et al. 2020). Furthermore, there is no cure for infected plants, and small WFT populations can infect many plants because a single insect can infect many plants over its adult life.

The capacity of WFT to transmit plant-infecting viruses can be severely deleterious, warranting the destruction (roguing) of plants in fields/greenhouses to prevent further spread of viral infection (Batuman et al. 2020). WFT transmit many viruses in the family *Tospoviridae*, a group for which TSWV is one of the most consequential plant viruses transmitted by thrips (Gilbertson et al. 2015). These virus infections can cause severe disease, depending on factors such as viral virulence and thrips population density (Batuman et al. 2020; Reitz et al. 2020). Like its thrips vector, TSWV has a wide host range among both crop and ornamental plants, constraining the production of peanuts, peppers, tobacco, and tomatoes, which causes annual economic losses in the hundreds of millions of dollars per crop (Pappu et al. 2009). Late second instar larvae (L2) or adult stage WFTs inoculate TSWV virus particles into healthy plant hosts, having previously acquired the virus as neonate larvae (L1) (Ullman 1992).

WFTs acquire viral particles in the L1 stage, during which the virus replicates in multiple organs, ultimately circulating to and invading the salivary glands, after which it is inoculated into hosts with saliva during feeding (Nagata et al. 2004; Montero-Astúa, Ullman, et al. 2016). TSWV accumulation (replication and transcription) in WFT is temporally dynamic, aligning robustly with thrips development, i.e., neometabolism (Ullman 1992; Montero-Astúa, Ullman, et al. 2016; Rotenberg & Whitfield 2018; Han & Rotenberg 2025). The highest virus titers occur in the late L2 stage, at a time when gut infection is extensive (Han & Rotenberg 2021) and the virus has breached the SGs before ecdysis into the propupal (P1) stage (Nagata et al. 2004; Montero-Astúa, Ullman, et al. 2016; Ullman 1992). Virus titers plummet precipitously during pupation, and then after adult eclosion, virus accumulates in the adult bodies (limited to SGs) but not to the level of the L2 stage (midgut and SG infection). The efficiency of virus inoculation by viruliferous adults declines with age (Rotenberg & Whitfield 2018; Reitz et al. 2020). Notably, while naïve adults can feed on TSWV-infected tissue to ingest and acquire virus, resulting in gut infection, they remain nonviruliferous due to a midgut barrier in adults that precludes viral particles from reaching the SGs (Ullman 1992; Montero-Astúa, Ullman, et al. 2016). Thus, virus entry of the late L2 SGs is a critical determinant of thrips vector competence.

Infection with TSWV has been shown to modify the feeding behavior of adult thrips. In adult males, relative to their female counterparts, infection with TSWV increased the frequency of important feeding behaviors, non-ingestion probes for example, that are related to inoculation efficiency (Stafford et al. 2011). Depletion of glycogen stores in adult infected thrips, after acquiring virus at the larval stages, is postulated as a possible explanation for the increased propensity to feed (Bailey, Kondragunta, Choi, Han, Rotenberg, et al. 2024; Bailey, Kondragunta, Choi, Han, McInnes, et al. 2024). Such apparent changes in insect physiology and behavior associated with virus infection have generated curiosity in the molecular factors underlying these effects (Han & Rotenberg 2025; Han & Rotenberg 2024).

Molecular events during TSWV infection within thrips are becoming increasingly clear as sequencing technologies advance (Schneweis et al. 2017; Stafford-Banks et al. 2014; Rotenberg et al. 2020b; Han & Rotenberg 2021). Because the salivary glands of thrips are critical to the insect’s ability to obtain nutrition and transmit viruses, there is considerable interest in identifying the relevant genes and proteins involved in SG processes during thrips growth and development (larval and adult) and virus infection. It has been reported that TSWV bi-directionally modulates expression of several genes that code for proteins involved in protein and complex carbohydrate catabolism, polysaccharide breakdown, calcium binding, detoxification, innate immunity, and several uncharacterized proteins in whole bodies (Schneweis et al. 2017; Zhang et al. 2013) and tissue systems, including adult SGs (Stafford-Banks et al. 2014; Han & Rotenberg 2021; Rajarapu et al. 2022). Using modern proteomic technologies that utilize small amounts of protein, analyses of adult WFT SG tissues revealed a sex-biased SG proteome-level response to TSWV infection, highlighting diverse cellular processes that may explain dimorphic feeding behavior and virus inoculation efficiency by adult thrips (Rajarapu et al. 2022).

The SGs of the WFT consist of a pair of principal and secondary glands each. The larger, ovoidal principal glands are composed of two lobes, each with eight, binucleated and loosely aggregated cells, whereas the secondary glands are tubular, flat and mononucleated (Ullman et al. 1989). A central duct connects the principal SG to the mouthparts and secretes primary saliva that together with secretions from the secondary SG are involved in tissue penetration, feeding, and virus transmission. Size, sex, and life-stage differences are associated with alterations in the salivary components of WFTs, which in turn are affected by virus infection status (Rajarapu et al. 2022). In this paper, we seek to clarify the connections between the TSWV, SG transcriptome, and the salivary proteome of WFT, in relation to sex and development to shed further light on how WFT interacts with TSWV and plant hosts. We reveal the extent to which TSWV affects the SG of larval (L2) and adult WFTs and the discordance between the transcriptome-level (present study) and proteome-level response (Rajarapu et al. 2022) to TSWV in adults. Our study revealed SG-specific transcriptomes, with the most profound effect of TSWV infection on the L2 stage. Our findings provide new knowledge about TSWV infection and associated cellular responses at the time the SG tissue barrier is first breached by the virus and open the door to molecular-based approaches to block transmission at the larval stage.

## Materials and Methods

### Thrips and virus inoculum sources

A western flower thrips (WFT) colony, originally collected from Hawaii and maintained in captivity since 1985, was reared on green bean pods (*Phaseolus vulgaris* L.) as described previously (Ullman 1992). Two-week old *Datura stramonium* plants, rub-inoculated with TSWV^MR-01^ were maintained in a greenhouse to serve as a source of inoculum for each experiment. Symptomatic plants, 8-10 days post rub-inoculation, were ideal for virus acquisition by first instar larvae. TSWV isolate MR-01 (TSWV ^MR-01^) was collected originally from infected radicchio in Monterey County, California and was flash frozen and stored at −80°C in the Ullman laboratory. Complete sequences of this strain can be found in the National Center for Biotechnology Information database (https://www.ncbi.nlm.nih.gov/nuccore), using accession numbers MG593199 (S RNA), MG593198 (M RNA), and MG593197 (L RNA).

### Virus acquisition and salivary gland dissections

For the transcriptomic experiments, salivary gland (SG) dissections were performed from three biological replicates each of non-infected and TSWV-infected second instar larvae (L2), adult females, and adult males. Biological replicates were obtained as follows: adult thrips were given 24 hours to lay eggs on green bean pods, after which they were removed. When hatching was first observed, the beans were gently brushed to remove any hatched thrips. These beans were transferred to a new container. After 12 hours, the newly eclosed larvae were placed in a container with new beans (to obtain non-infected L2s) or onto TSWV-infected *Datura stramonium* L. leaves for 12-16 hours, after which they were given green bean pods (to obtain infected L2s). An additional 72 hours after placement onto either green bean pods or TSWV-infected leaves, the L2s were collected for dissections (72-84 h-post egg eclosion). Adult females and males were collected approximately 72 hours after molting to adulthood from either noninfected or infected cohorts. Each biological replicate consisted of pooled SGs surgically removed from a total of 150 individuals that were collected 25 at a time at each dissection event. The resulting biological replicates consisted of non-infected second instar larvae (NLSG), non-infected females (NFSG), and non-infected males (NMSG) and their respective infected cohorts (ILSG, IFSG, and IMSG). To identify SG-enriched genes, previously available whole-body RNA-seq datasets were used for adult samples (Bailey et al., 2024b) and larval samples (Schneweis et al. 2017). During dissection, SGs were obtained from each noninfected and infected thrips life stage described above. To minimize cross-contamination, dissection tools were dedicated to each biological group. Teflon-coated blades were used to decapitate thrips behind their forelegs while placed in 25 µL of chilled 50% ethanol deposited on a cold microscope slide. The SGs were separated from other gut contents using a single hairbrush. All dissections were conducted within a consistent 4-hour window to control for any differences due to time of day.

### RNA extraction and preparation

Total RNA was extracted from the respective pooled SG samples using TRIzol according to the manufacturer’s recommendations. In brief, microcentrifuge tubes containing frozen SGs in 100 µL TRIzol were removed from the −80 ⁰C freezer and ground up using a tightly fitting pestle. An additional 400 µL of TRIzol was used to wash off the thrips lysate into the tube. Incubation at room temperature for 5 minutes was followed by adding 100 µL of chloroform to each tube, mixing, and incubating again for another 3 minutes. The samples were then spun at 12,000 x g for 15 minutes at 4 ⁰C. The resulting aqueous phase was carefully transferred into a new tube with a pipette.

Due to the small amount of starting sample, 5 µg of RNase-free glycogen was added to each tube as a carrier with 250 µL of isopropanol. To ensure high yield of resulting RNA extract, an overnight incubation at −20 ⁰C was carried out. Following, the tubes were centrifuged at 12,000 x g for 10 minutes at 4 ⁰C to precipitate RNA. Total RNA was resuspended in 20 µL of nuclease-free water, and contaminating DNA was removed with the TURBO DNA-free kit from Invitrogen.

Poly(A) libraries were prepared to increase enrichment for mRNA with a TruSeq RNA Library Prep Kit (Illumina, San Diego, CA, USA) and sequenced by the DNA Sequencing and Genotyping Core at the Cincinnati Children’s Hospital Medical Center (CCHMC). RNA was quantified using a Qubit 3.0 Fluorometer (Life Technologies, Carlsbad, CA), and RNA integrity was assessed using an Agilent Bioanalyzer (Santa Clara, CA, USA). All samples had an RNA Integrity Number (RIN) over 7. Total RNA (150–500 ng) was poly(A)-selected and reverse-transcribed using the TruSeq Stranded mRNA Library Preparation Kit (Illumina). Each sample was fitted with a sample-specific barcode for multiplexing. Following 15 cycles of polymerase chain reaction amplification, the completed libraries were sequenced on a HiSeq 2500 (Illumina) in Rapid Mode.

### RNAseq analyses of thrips salivary glands

Bioinformatic analyses were conducted based on our previous methods for analyses in thrips (Rotenberg et al. 2020b; Bailey, Kondragunta, Choi, Han, Rotenberg, et al. 2024). Importantly, we used two independent pipelines to analyze the RNA-seq data (Davies et al. 2021; Pathak et al. 2022). In CLC Genomics Workbench version 12.0 (QIAGEN), the quality of the reads was evaluated before and after trimming with FastQC (Andrews & Others 2010). The trimmed/cleaned, paired-end reads were mapped to the official gene set from the *Frankliniella occidentalis* genome (GenBank assembly GCA_000697945.4, Focc_3,(Rotenberg et al. 2020b)). The mapped reads were normalized to transcripts per kilobase per million (TPM) as a proxy for expression. EDGE methods with default settings in CLC Genomics Workbench 12 (QIAGEN) with false discovery rate (FDR) corrected p-values less than 0.05 were considered significant. For the second pipeline, an RNA-seq analysis protocol was developed in the Galaxy system (Afgan et al. 2018). The datasets were trimmed for adapters and cleaned using Trimmomatic (Bolger et al. 2014). The reads were mapped to the predicted gene set and quantified using Sailfish (Patro et al. 2014) to generate normalized expression values as TPM. Mapped read counts were summarized and combined using tximport (version 0.1). Differential expression was analyzed using DESeq2 (Love et al., 2014), and false discovery rate (FDR)-adjusted p-values < 0.05 were considered significant. For the SG-enrichment analyses, a specific transcript was considered to be enriched in the SG if the expression was 4-fold higher in the SG compared to the whole body. A Pearson correlation coefficient was calculated to assess overlap between differentially expressed transcripts from each pipeline; it was 0.94. Due to the high consistency, subsequent analyses focused on the second pipeline. Data sets with differential expression were examined for enriched Gene Ontology categories using g:Profiler (Raudvere et al. 2019) and summarized with REVIGO (Supek et al. 2011). GO enrichment results were summarized and visualized using the rrvgo R package (Sayols 2023). Pairwise semantic similarity among enriched GO terms was calculated, and highly redundant terms were clustered based on semantic similarity to reduce redundancy while retaining representative biological processes. The resulting representative GO terms were visualized to facilitate biological interpretation of the enriched functional categories. RNA-seq data sets are available through the NCBI SRA under the following bioprojects: PRJNA1311131 (salivary glands), PRJNA454326 (whole larvae), and PRJNA852191 (whole adults).

In addition to assessing expression changes in thrips, the presence of TSWV was confirmed in the salivary glands by detecting viral transcripts. These methods were based on those previously established to identify viral presence in insect RNA-seq datasets (Wu et al. 2020; Liu et al. 2011; Britt-Ugartemendia et al. 2025). To do so, reads from each sample were mapped to the TSWV genome (segment S - MG593199.1; segment M - MG593198.1; segment L - MG593197.1) using Salmon (Patro et al. 2017) and Kallisto (Bray et al. 2016) with baseline parameters. The presence of the virus was determined by mapping viral reads to each genome segment; at least 10 reads were detected across all three segments, and quantification was obtained using both Kallisto and Salmon. Tables S1-S3 confirm infection status based on detected viral reads.

### Weighted gene co-expression network analysis (WGCNA)

We performed a comparative analysis of each set and constructed a gene co-expression network using the WGCNA R software package (https://horvath.genetics.ucla.edu/html/CoexpressionNetwork/Rpackages/WGCNA/). This method enabled us to identify genes with similar expression patterns in the salivary glands across developmental stages and sexes, group them into modules, and associate them with the corresponding samples. To prepare the expression data for WGCNA, we removed genes with zero variance (e.g., genes with expression values of 0 across all sets) and included genes with a minimal expression level of 10 TPM across all samples, resulting in a dataset of 10,217 genes to construct an unsigned network. We selected a soft thresholding power of 12 based on the scale-free topology fit index curve, which was generated before network construction. The minimum module size was set to 20. To associate each module with specific developmental stages, the developmental stages were included as input traits during network construction. Modules that showed significant correlations with the traits (p < 0.05) were further examined to explore their functions and relationships with developmental stages and sexes.

### Comparison of salivary gland transcriptome and proteome

The total salivary gland proteome, including 2663 proteins from infected and non-infected adults (Rajarapu et al. 2022), were used to identify the transcripts encoding these proteins in the salivary gland transcriptome of the current study. Normalized count data of the transcripts and the protein abundances were log2 transformed to generate correlation curves between the proteins and the transcripts encoding the corresponding transcripts in JMP Pro 16 software. Transcripts with a cut-off of −2 or less were excluded from the correlation analysis, as they showed no correlation between protein and transcript abundances. The significance of correlation was determined by Spearman analysis.

## Results & Discussion

### Genes with enriched expression in the salivary glands

Previous studies have established salivary gland-enriched gene sets (Rotenberg et al. 2020a) using pooled male or female RNAseq datasets (+/- TSWV) available at the time. In the present study, our investigation revealed distinct differences in expression between salivary gland and whole-body samples of non-exposed thrips (Fig. 1A-C). Of the predicted genes, 914 (Table S1), 2747 (Table S2), and 1580 genes were enriched in females, males, and larvae, respectively (Fig. 1D). Of these, 357 were highly enriched in all stages (overlapping set of genes), likely representing the core transcriptional factors of thrips SGs (Fig. 1E). Of these 357, 44 have predicted signal peptides, suggesting these could be associated with the secreted saliva (Table S4-S6). GO analysis of this overlapping set revealed enrichment in many processes associated with tissue growth and development (Fig. 2). Specific GO analysis for each set is included in Figure S1. These initial studies provide an extensive transcriptional profile of the SG of thrips.

**Figure 1.**
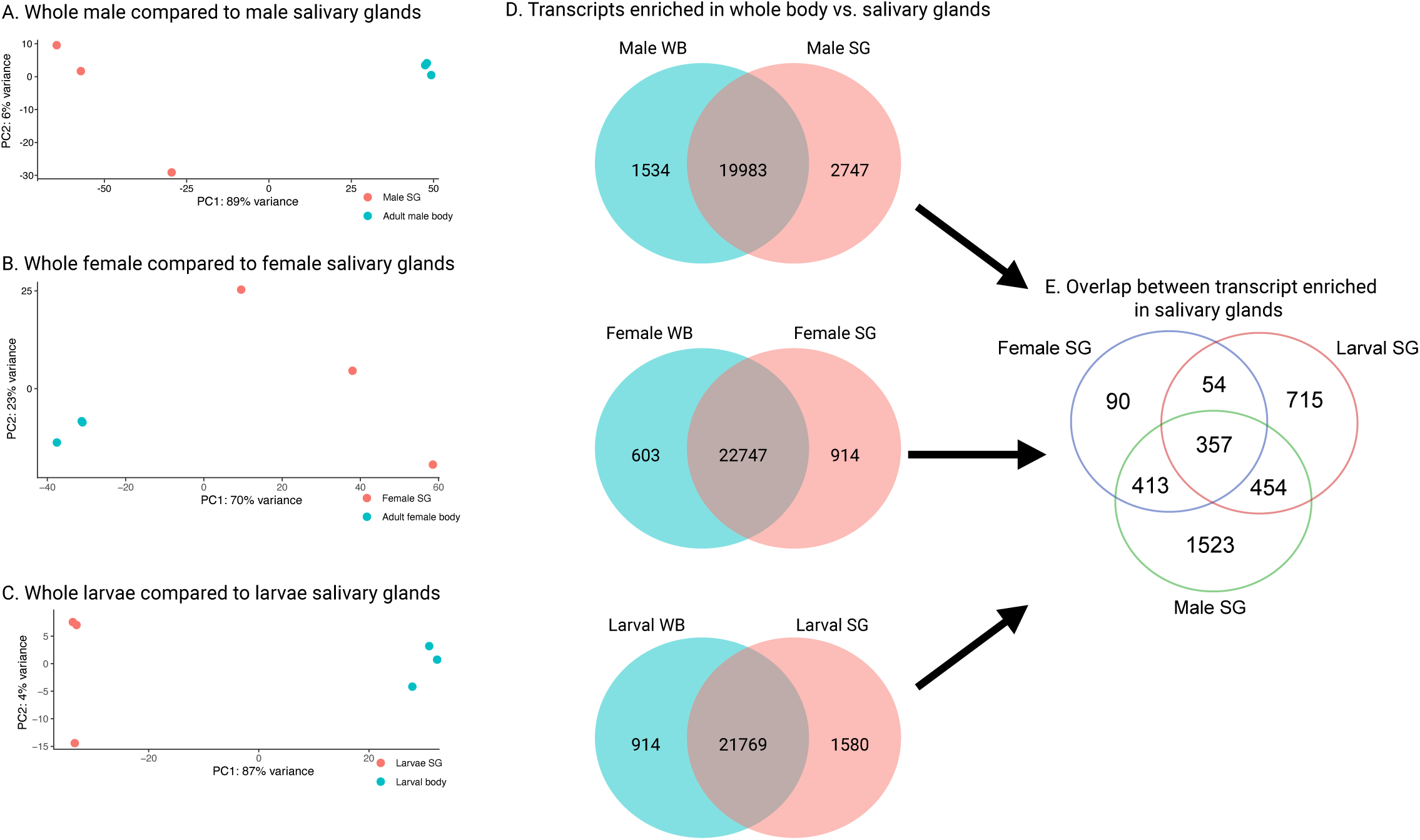
Enrichment of transcript-level expression in male, female, and second instar larval salivary glands of *Frankliniella occidentals* in relation to whole bodies. (A–C) Principal component analysis (PCA) of normalized gene expression profiles comparing salivary gland (SG) samples with corresponding whole-body (WB) samples from adult males (A), adult females (B), and larvae (C). Biological replicates cluster according to tissue type, demonstrating that SGs exhibit transcriptional profiles distinct from those of whole-body samples. (D) Venn diagrams showing the number of transcripts with significantly enriched expression in either the SG or WB for males, females, and larvae (false discovery rate-adjusted *P* < 0.05). The overlap represents transcripts shared between SG and WB, whereas the non-overlapping regions represent tissue-enriched transcripts. (E) Venn diagram illustrating the overlap among SG-enriched transcript sets from males, females, and larvae, highlighting transcripts shared across all three groups as well as those uniquely enriched in each developmental stage or sex, indicating both conserved and stage- or sex-specific components of the salivary gland transcriptome.

**Figure 2.**
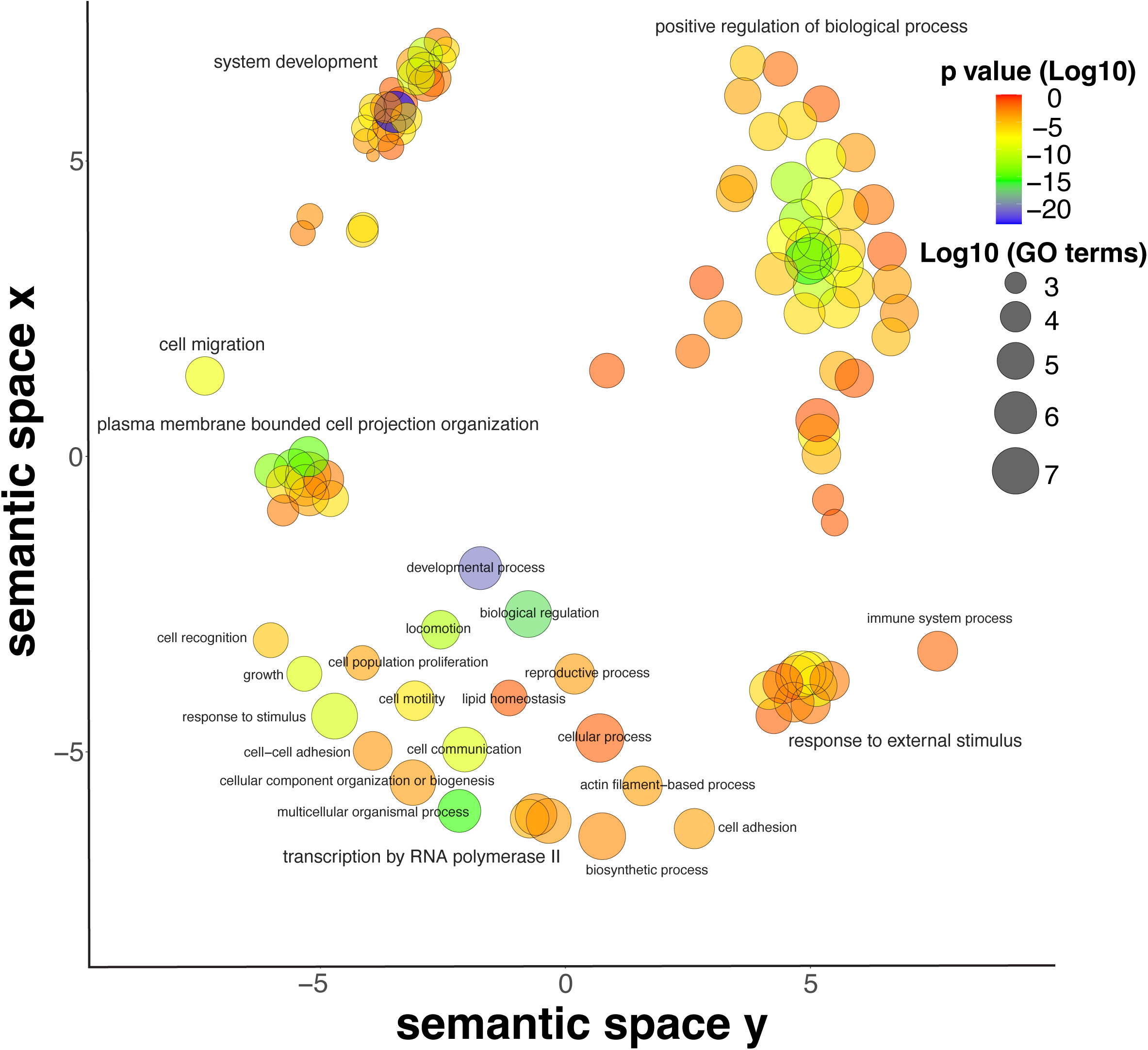
Gene ontologies provisionally assigned to the core set of transcripts with enriched expression in the salivary gland of *Frankliniella occidentalis*. These analyses focus on the 357 SG-associated genes (Fig. 1E) shared among males, females, and second-instar larval thrips. Gene ontology (GO) terms were established with g:Profiler (Raudvere et al. 2019), which clusters GO terms according to semantic (i.e., meaning) similarity and projects them into a two-dimensional semantic space. Each circle represents a GO Biological Process term, with the distance between circles reflecting their semantic relatedness. Circle size is proportional to the frequency of the GO term within the GO database, with larger circles representing more general terms and smaller circles representing more specific terms. Circle color indicates the significance of GO enrichment (−log_10_ p-value), with warmer colors corresponding to more highly significant terms. Labels identify representative GO terms for major functional clusters, illustrating enrichment of biological processes.

A few studies have examined the composition of *F. occidentalis* salivary glands (Rotenberg et al. 2020a; Stafford-Banks et al. 2014; Rajarapu et al. 2022). In one study, an adult-specific comparison of male and female whole bodies and salivary glands revealed 123 transcripts enriched in expression in the SGs, including lipases, mannanases, and deoxyribonucleases, as well as many uncharacterized transcripts and novel *F. occidentalis* proteins (Rotenberg et al. 2020a). That study was limited to single RNA-seq samples per sex (albeit validated by replicated qPCR data), and therefore many genes (transcripts) with more muted differences in expression were likely not represented in the SG-enriched dataset. Here, we show substantial differences in SG transcriptomes among males, females, and larvae, extending to differences in the GO categories enriched in the SG at each stage (Fig. S1), with only 40% of those previously identified observed in our analysis. A similar difference was observed in tsetse fly salivary components, where expanded in-depth transcriptome analyses revealed a more extensive set of SG-associated genes (Telleria et al. 2014; Alves-Silva et al. 2010; Matetovici et al. 2016; Attardo et al. 2019). In the present study, we expected differences in SG expression between males and females, as this was the case at the proteome level (Rajarapu et al., 2022). The wide differences between the larval and adult stages were partly unexpected, as all stages feed on the same food source, although foliage microhabitats vary. After hatching from eggs, larvae of other thrips species have been observed to predominantly feed on the underside of leaves and newly emerged foliage (Varga et al. 2010). The adults feed on other, more mature foliage, flower parts and pollen, suggesting a divergence in feeding between the stages of this pest. Also, these differences could be due to developmental shifts in behavior or morphological differences in the mouthparts, such as size differences, or could reflect differences in subsequent digestive processes that require different preprocessing by the saliva.

Several genes among those that are enriched are consistent with functions in extra-oral digestion during plant feeding (Cantón & Bonning 2020; Stafford-Banks et al. 2014), including trypsin-1-like and aminopeptidase N-like proteases, which are likely involved in the hydrolysis of plant proteins, and phospholipase A1-like, which may facilitate disruption of plant cell membranes to release cellular contents (Fig. 1, Table S4-S6). Multiple carboxylesterases may contribute to the metabolism of dietary lipids and detoxification of plant defensive compounds encountered during feeding (Vogel et al. 2014). In addition, abundant salivary proteins such as mucins and proline-rich proteins, together with glycosylation enzymes including polypeptide N-acetylgalactosaminyltransferases, suggest extensive production and post-translational modification of secreted salivary proteins (He et al. 2024; Ma et al. 2024; De Schutter 2026). Notably, relatively few classical plant cell wall-degrading enzymes were identified (Tran et al. 2024), supporting the view that thrips rely primarily on mechanical puncture and disruption of cells (Ullman et al. 1992; Chisholm & Lewis 1984), combined with proteolysis and membrane disruption, rather than on extensive degradation of structural polysaccharides by the saliva, to access nutrients from plant tissues.

### Impact of TSWV infection on salivary gland transcriptome

Detectable levels of TSWV transcripts were observed in all RNA-seq sets from thrips that had previously fed on infected plant material (Table S1-3). When the transcriptional response was examined in relation to TSWV infection, there were differences between the adult sexes and larvae (Fig. 3A). TSWV infection of larval SG resulted in the greatest number of differentially expressed genes, with significant differences in expression of 1.1% and 2.4% of the L2 SG transcriptome in the noninfected and infected treatments, respectively (Fig. 3B), compared to 0.26% (control) and 0.34% (infected) of the male SG transcriptome, and approximately 0.13% of the female SG transcriptome, irrespective of infection status. Like what was previously observed in the proteome (Rajarapu et al. 2022); there was little to no overlap among the differentially abundant genes responsive to TSWV across males, females, and larvae (Fig. 3C), suggesting that SG responses differ drastically across stages. This finding is consistent with previously reported developmental stage-specific responses to TSWV infection of *Frankliniella* tissues (Schneweis et al. 2017; Shrestha et al. 2017; Han & Rotenberg 2021; Rajarapu et al. 2022) and may point to the tight link between thrips development and tissue tropism, i.e., spatiotemporal dissemination of TSWV in the thrips vector body (reviewed in (Montero-Astúa, Stafford-Banks, et al. 2016)). When only SG-enriched transcripts (compared to whole bodies) were examined in relation to infection, larvae were more profoundly affected than adults (Fig. 3). As reported in SG proteomes and transcriptomes of closely related species (Rotenberg et al. 2020b), our study also revealed numerous uncharacterized and hypothetical proteins (Fig. 4), which represented 10% of the larval transcriptome shifted during infection.

**Figure 3.**
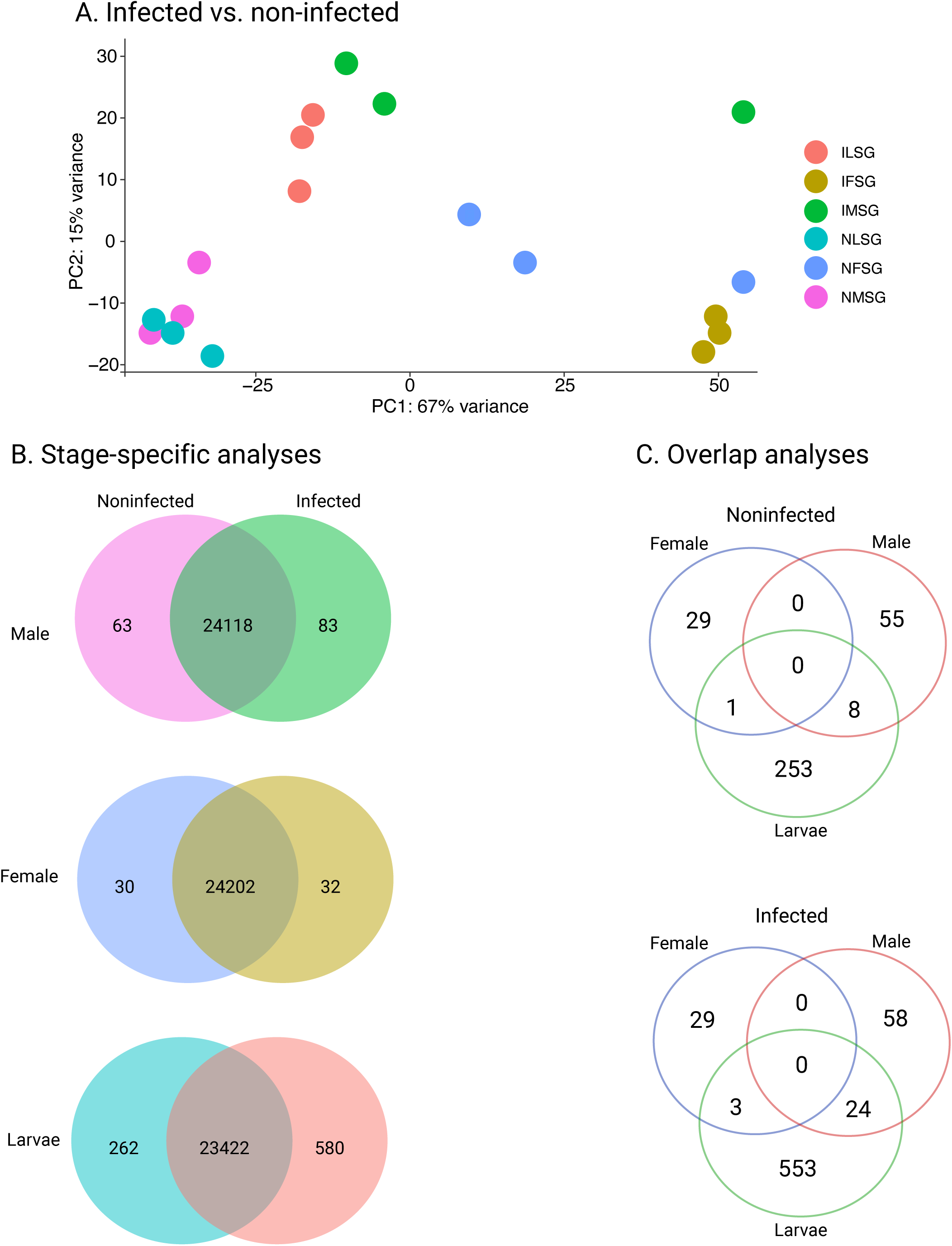
Shifts in transcriptome-level expression in salivary glands from second instar larvae, females, and males of *Frankliniella occidentalis* infected with tomato spotted wilt virus (TSWV). A) Principal component analysis (PCA) of normalized gene expression profiles from salivary glands (SGs) of TSWV-infected (I) and noninfected (N) larvae (L), adult females (F), and adult males (M). Samples cluster according to developmental stage, sex, and infection status, demonstrating that TSWV infection alters the SG transcriptome while maintaining distinct stage- and sex-specific expression patterns. (B) Venn diagrams comparing the number of transcript sequences expressed between noninfected and TSWV-infected SGs within each developmental stage or sex. Differentially-expressed transcripts were identified using a false discovery rate (FDR)-adjusted significance threshold (*P* < 0.05), with overlapping regions representing transcripts expressed in both conditions and non-overlapping regions representing transcripts whose expression was specific to either infected or noninfected SGs. (C) Venn diagrams illustrating the overlap of transcripts differentially expressed in response to TSWV infection among larvae, adult females, and adult males. Separate analyses are shown for transcripts enriched in noninfected and infected SGs, highlighting both conserved and stage- or sex-specific transcriptional responses to viral infection.

**Figure 4.**
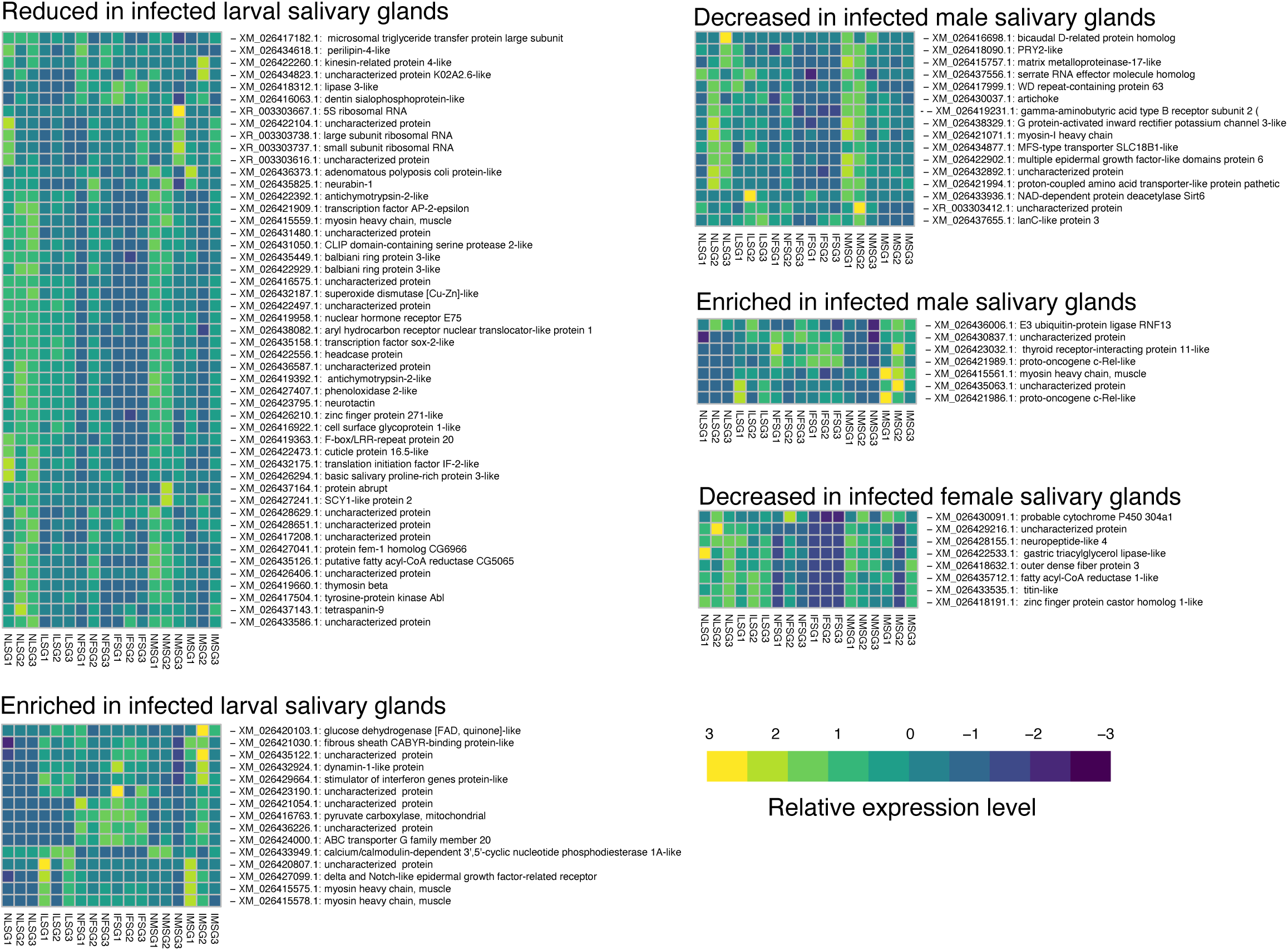
Expression levels of genes differentially expressed following infection with tomato spotted wilt virus (TSWV) in second instar larval and adult salivary glands of *Frankliniella occidentalis*. Heatmaps showing the relative expression of transcripts that were significantly differentially expressed between TSWV-infected and noninfected salivary glands (SGs) of larvae, adult females, and adult males (false discovery rate-adjusted *P* < 0.05) and enriched in SGs. Transcripts are grouped according to those exhibiting increased or decreased levels in response to infection within each developmental stage or sex. Columns represent individual biological replicates of noninfected larval SGs (NLSG), infected larval SGs (ILSG), noninfected female SGs (NFSG), infected female SGs (IFSG), noninfected male SGs (NMSG), and infected male SGs (IMSG), while rows represent individual transcripts annotated by their predicted gene products. Gene expression values are displayed as normalized, row-scaled expression levels (Z-scores), with warmer colors indicating relatively higher transcript abundance and cooler colors indicating relatively lower transcript abundance across samples. Note that there was no significant group of enriched, differentially-abundant transcripts in infected female SGs.

Gene ontologies for all differentially expressed genes in L2 SGs (842; Figure 3) revealed numerous altered processes, with significant modulation in growth-related aspects, suggesting that developmental processes in the L2 SG tissue may be affected by viral infection (Fig. 5, Fig. S2). The combined gene set was enriched for biological processes including anatomical structure development, cellular anatomical entity morphogenesis, cellular component biogenesis and organization, developmental and metabolic processes, reproductive processes, and ribosome biogenesis (Fig. 5).

**Figure 5.**
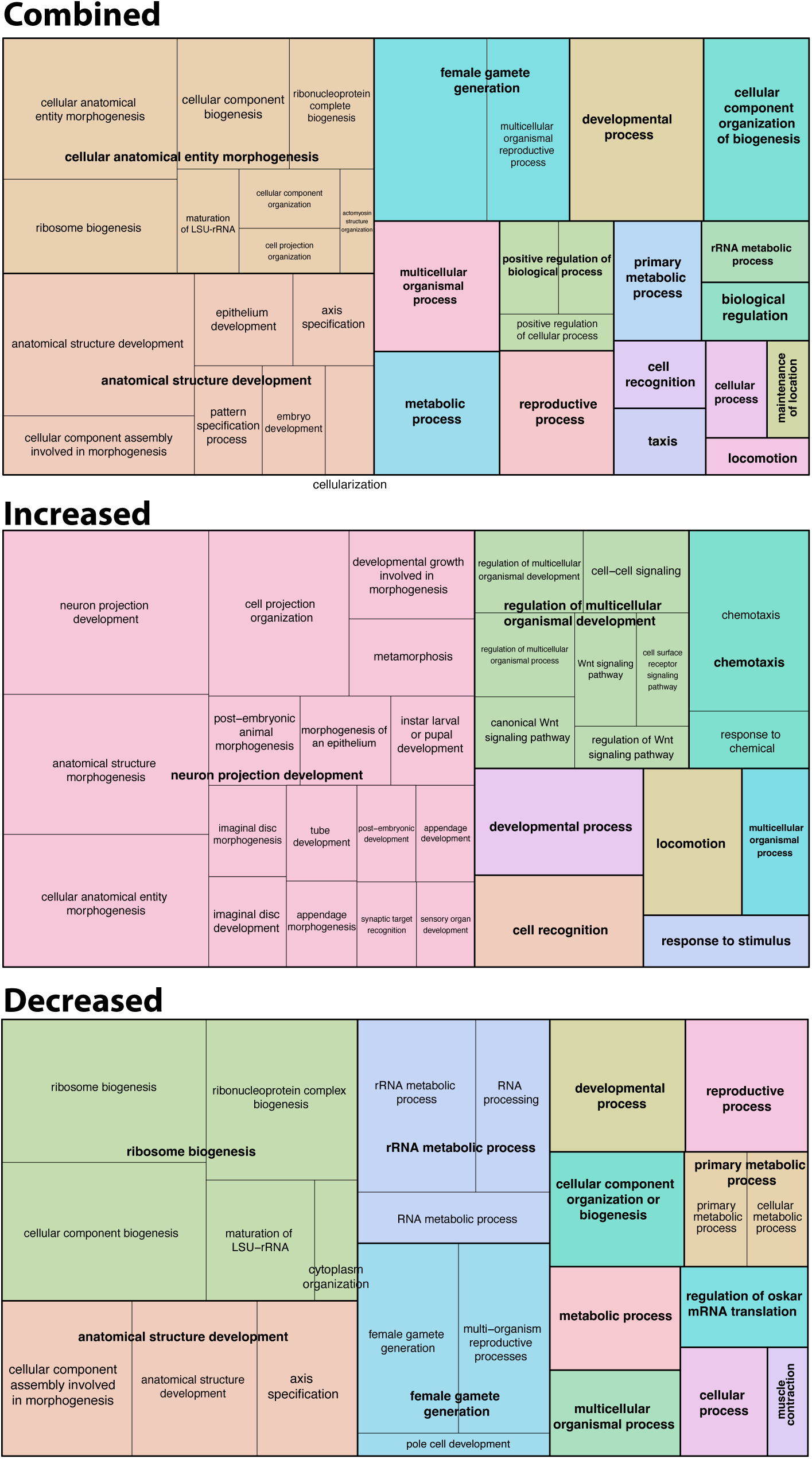
Gene ontologies associated with genes differentially expressed in the salivary gland of second instar larvae of *Frankliniella occidentalis* infected with tomato spotted wilt virus (TSWV). Gene ontology (GO) enrichment analysis was performed on transcripts that were differentially expressed in TSWV-infected larval salivary glands relative to noninfected larvae. Expressional differences are based on Table S11 and S12. Gene ontology terms were established with g:Profiler (Raudvere et al. 2019) based on larval transcriptome. Specific GO terms were combined with REVIGO to generate a treemap (Supek et al. 2011). Treemaps illustrate GO categories for the combined set of differentially expressed transcripts as well as transcripts exhibiting increased or decreased expression following TSWV infection. Each rectangle represents an enriched GO term, with related terms grouped into larger semantic clusters, and the area of each rectangle reflects the relative significance of the associated GO term.

Upregulated genes were primarily associated with neuron projection development, anatomical structure morphogenesis, cell projection organization, canonical Wnt signaling, cell–cell signaling, chemotaxis, and responses to chemical stimuli, suggesting activation of developmental remodeling and signaling pathways during infection. In contrast, downregulated genes were enriched for ribosome biogenesis, ribonucleoprotein complex biogenesis, rRNA metabolic processes, RNA processing, and cellular component biogenesis, as well as developmental, metabolic, and reproductive processes, indicating suppression of protein synthesis and RNA metabolism in response to salivary gland infection (Fig. 5).

The GO enrichment analysis indicates that salivary gland infection triggers extensive transcriptional reprogramming in larvae, characterized by the activation of developmental, morphogenetic, and signaling pathways, while simultaneously suppressing ribosome biogenesis, RNA metabolism, and protein synthesis. These changes suggest that infected tissues prioritize cellular remodeling and immune-related responses over growth and biosynthetic activities, reflecting an adaptive response to infection. Importantly, TSWV uses host mRNA cap snatching for its replication (Duijsings et al. 2001), which could significantly compete with the transcriptional and translational processes of the cell (Duijsings et al. 2001), as suggested by suppressed RNA biogenesis, metabolism and protein synthesis. WGCNA on the gene expression data (normalized abundance of each transcript) identified specific modules of genes with similar expression for each developmental stage and sex, however, infection status alone (‘infected’ or ‘non-infected’) had no apparent correlation with the WGCNA module of co-expressed genes (Fig. 6). Like the proteomic responses reported in Rajarapu et al. (2022), our WGCNA analysis was primarily explained by sex/stage differences in *F. occidentalis*. Both studies provide a thorough description of SG-associated factors for *F. occidentalis* in relation to TSWV.

**Figure 6.**
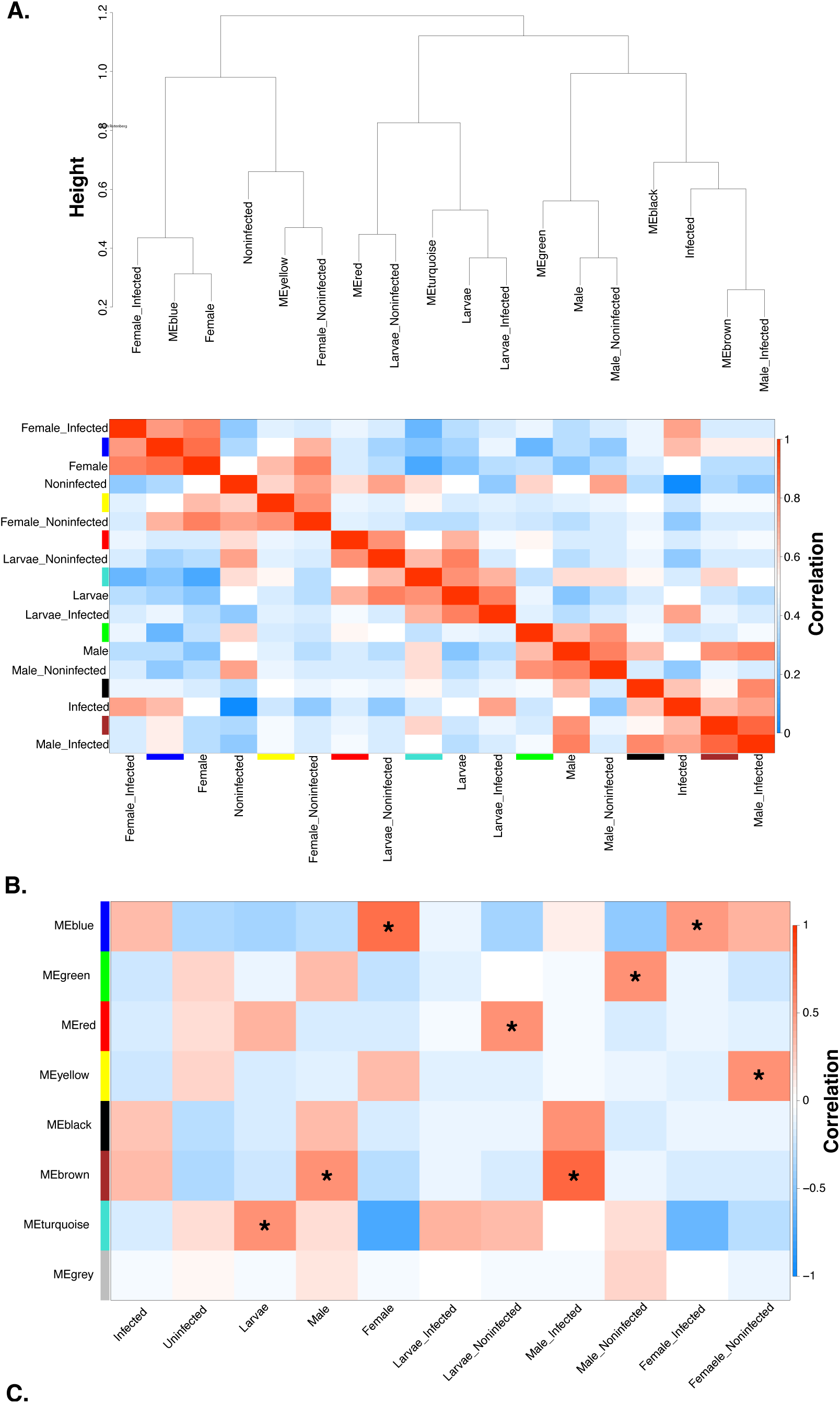
Weighted gene co-expression network analysis (WGCNA) for second instar larvae and adult salivary glands of *Frankliniella occidentalis*. (A) Hierarchical clustering dendrogram illustrating the relationships among module eigengenes and experimental traits. Module eigengenes, representing the first principal component of each co-expression module, were clustered together with experimental traits to evaluate similarities in global expression patterns. Traits associated with infection status (infected and noninfected), developmental stage (larval), sex (male and female), and stage- or sex-specific infection groups cluster according to the similarity of their eigengene expression profiles, revealing relationships among transcriptional modules and biological conditions. Branch lengths indicate the degree of dissimilarity between module eigengenes and experimental traits, with shorter branch lengths reflecting greater similarity. (B) Heatmap showing correlation coefficients between module eigengenes and experimental traits, including infection status, developmental stage, sex, and the corresponding stage- and sex-specific treatment groups. Each row represents a co-expression module (identified by module color), and each column represents an experimental trait. Cell color indicates the strength and direction of the correlation, with red denoting positive correlations and blue denoting negative correlations. Asterisks (*) indicate statistically significant module-trait associations (*P* < 0.05), identifying co-expression modules that are significantly associated with specific biological conditions.

Many individual transcripts increased during TSWV infection and encode uncharacterized proteins and non-coding RNAs, suggesting that virus infection suppresses a diverse range of host genes whose functions remain largely unknown (Fig. 5, Table S11). Among the annotated genes, transcripts associated with lipid metabolism (e.g., lipase-like proteins), amino acid metabolism (pyrroline-5-carboxylate reductase), extracellular matrix-related proteins (elastin-like), and developmental regulators (maternal effect lethal 26-like protein) exhibited reduced expression. Additionally, the downregulation of regulatory non-coding RNAs may indicate disruption of host gene regulatory networks during larval infection. Several genes were significantly more abundant in the salivary glands of noninfected larvae. In addition to numerous uncharacterized non-coding RNAs, transcripts encoding large and small subunit ribosomal RNAs and 5S ribosomal RNA were significantly decreased in infected larvae, suggesting suppression of the host translational machinery. Several metabolic and digestive genes, including lipase 1-like, endothelial lipase-like, trypsin-7-like, trypsin alpha-like, and cathepsin L1-like, also exhibited lower expression during infection, suggesting altered digestive and lipid metabolic processes from saliva produced by infected salivary glands that could prompt the increased feeding observed during TSWV infection (Bailey, Kondragunta, Choi, Han, McInnes, et al. 2024; Stafford et al. 2011). Furthermore, genes involved in detoxification and cellular homeostasis, such as cytochrome P450 6K1-like, cytochrome P450 4C3-like, and cytochrome b5-related protein-like, were downregulated, suggesting that viral infection modifies xenobiotic pathways (Stavropoulou et al. 2018; Lu et al. 2021). Reduced expression of low-density lipoprotein receptor-like, proton-coupled amino acid transporter 4-like, and glucose transporter type 1 further indicates disruption of nutrient transport and cellular metabolism with TSWV infection in larval SG. Collectively, these findings suggest that viral infection suppresses genes involved in protein synthesis, digestion, detoxification, lipid metabolism, and nutrient transport, reflecting extensive reprogramming of salivary gland physiology that may favor viral replication and transmission during TSWV infection of larval SG tissues.

The present study uniquely provided new insights into the thrips-virus interaction during the developmental stage (L2), when the virus first breaches the SGs (Wijkamp & Peters 1993; Montero-Astúa, Stafford-Banks, et al. 2016), along its dissemination route from the gut to the principal SGs. As evidenced by the number of TSWV-responsive genes in the L2 compared to adult SGs, this breach and subsequent infection of the larval SGs had a substantial impact on SG response. Males showed the highest absolute fold changes in expression levels in their SGs during TSWV infection, which supports previous observations of a proteome shift (Rajarapu et al. 2022), but these differences were not as substantial (less than 1-fold differences among males, females, and larvae). The muted response of adult SGs to TSWV infection compared to L2 SGs may reflect the recency of infection of this tissue and prolonged tolerance to infection, which is a common aspect associated with prolonged viral infections (Kane & Golovkina 2010; Castelló-Sanjuán et al. 2025). This differs from observations in whole thrips bodies (Schneweis et al. 2017; Shrestha et al. 2017), where the virus is likely propagating in other organs and could affect processes such as reproduction. The processes inferred by the SG-enriched, differentially-abundant transcripts in infected L2s SGs paint a picture of a shift in many cellular processes. Notably, factors involved in anatomical development and ribosome biogenesis appeared suppressed, suggesting reduced salivary gland development and saliva protein production. The suppression of these factors is suggestive of “host shut-off” that occurs during infection with cap snatching viruses, such as TSWV. Cap snatching, which occurs in all orthotospoviruses and many other viruses, occurs when viral enzymes cleave and remove the entire cap structure from the 5’ end of host mRNA. The virus then uses this stolen cap leader to prime the synthesis of new viral mRNAs that start with a host-derived cap. The cell’s ribosomes then translate viral proteins. As host mRNAs are cap-depleted, translation of host proteins decreases. This process results in a shutdown of host cell protein production, which could, in part, explain the transcriptome-level response we observed in this study. Overall, TSWV-infected larvae exhibit distinct differences in processes associated with tissue growth and development, suggesting that SG function may be impaired following TSWV infection, thereby increasing feeding, as observed at other stages (Stafford et al. 2011). Of particular interest are increases in factors that underlie chemotaxis, locomotion, and neuronal function, suggesting that viral infection may affect thrips behavior. These neuronal changes could underscore the behavioral shifts previously observed in TSWV-infected WFT (Bailey, Kondragunta, Choi, Han, McInnes, et al. 2024).

### Overlap between transcriptome and proteome analyses

A previous study identified proteome shifts associated with thrips salivary glands in WFT males and females in relation to sex and TSWV infection status (Rajarapu et al. 2022). Our transcriptomic analyses identified 5012 salivary gland transcripts in the adults that overlap with 2663 proteins identified in the adult SG proteome (Rajarapu et al. 2022), revealing that these approaches have a significant but weak correlation overall in TSWV-infected and non-infected SGs (Fig. 7). These correlation patterns remain consistent when females and males are analyzed separately (Fig. 7). This is not surprising as weak correlations between protein and cognate transcript levels are common, due to the discordant temporal regulation of protein and transcript abundance in the cell. This significant but weak correlation between protein and cognate transcript abundance is consistent with findings for gut tissue of *F. occidentalis* (Han and Rotenberg, 2025); however, infection with TSWV tightened the correlation in that study. Most of the transcripts that contributed to the weak correlation in the present study encode cytoskeletal proteins. Between the proteome and the transcriptome, only a single transcript/protein was significantly modulated in abundance by TSWV in both data sets, which was integrin-P-bS. Cytoskeletal proteins are commonly associated with viral infection in insect systems (Khorramnejad et al. 2021; Horníková et al. 2022; Cui et al. 2020), and have been observed in other thrips species (Shrestha et al. 2017). Integrin-b-PS have been documented to be associated with the interaction between viruses and cells (Kausar et al. 2022; Li et al. 2007). Importantly, integrins are involved in the immune response of insects (Nainu et al. 2015); thus, integrin-b-PS could be involved in the thrips antiviral immune response. In addition to their role in infection, integrins have been associated with SG development and function (Jattani et al., 2009; Pirraglia et al., 2013), suggesting that the dynamics between thrips and TSWV within SGs could directly affect SG development.

**Figure 7.**
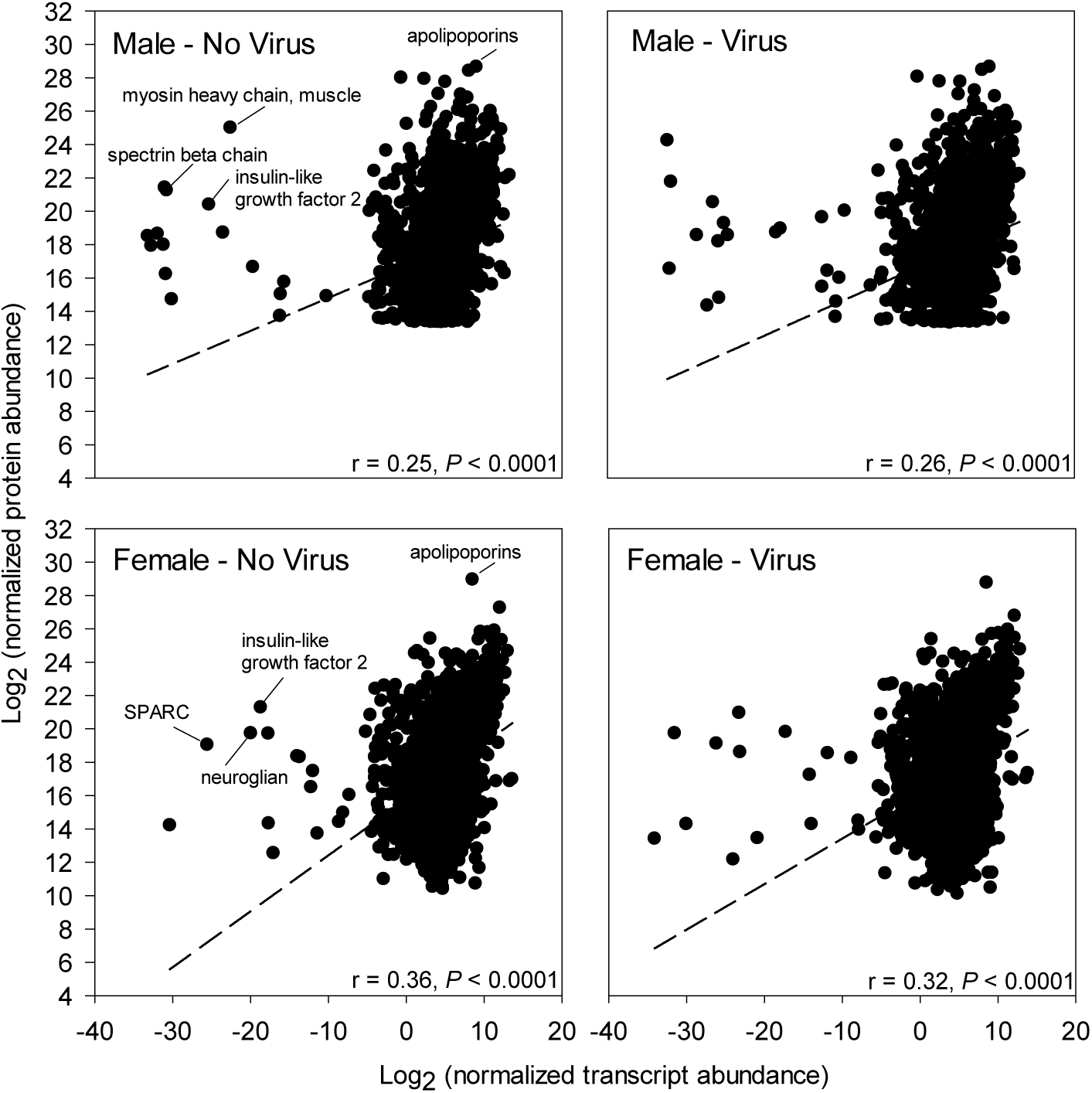
Comparative analysis between normalized abundance of proteins and their cognate transcripts in adult salivary glands of *Frankliniella occidentalis* infected with tomato spotted wilt virus (TSWV). Scatterplots comparing normalized transcript abundance (RNA-seq) and normalized protein abundance for salivary gland-associated genes in adult male and female thrips under noninfected and TSWV-infected conditions. Proteomic data were obtained from the previously published salivary gland proteomes of adult males and females (Rajarapu et al. 2022) and integrated with the transcriptomic datasets generated in the present study. Top panels compare transcript and protein abundance in noninfected (left) and TSWV-infected (right) males, while bottom panels show corresponding comparisons for noninfected (left) and TSWV-infected (right) females. Each point represents an individual sequence detected in both the transcriptomic and proteomic datasets. Dashed lines indicate the best-fit linear relationship between transcript and protein abundance. Pearson correlation analyses revealed weak but statistically significant positive correlations between transcript and protein abundance under all four conditions, indicating that transcript levels explain only a portion of the observed variation in protein abundance, while demonstrating an overall concordance between the two datasets. A few extreme datapoints, identified in all four conditions, are indicated by their gene description for the noninfected condition. SPARC = secreted protein, acidic and rich in cysteine); structural protein associated with the extracellular matrix.

## Concluding remarks

These studies provide the complete transcriptional characterization of viral infection in the WFT salivary glands. Three primary take-home results emerge: significant differences in SG-enriched gene sets among larvae, males, and females suggest underlying, unknown differences in feeding between sexes and developmental stages. Importantly, this has not been studied in detail or WFT, where most feeding studies have assumed that larvae and adults are feeding similarly (Reitz et al. 2020; Schneweis et al. 2017; Rotenberg et al. 2020b). The differences observed between sexes and developmental stages highlight the need to examine feeding strategies in greater detail. As an example, male mosquitoes, which feed only on sugar, and female mosquitoes, which feed on sugar and blood, show similar levels of differences to those we observed in this study (Ribeiro et al. 2016).

Alternatively, thrips larvae and adults differ in their digestive physiology, reflecting their distinct developmental stages and nutritional demands. In general, larvae exhibit higher digestive and metabolic activity to support rapid growth and development, whereas adults allocate more resources toward maintenance and reproduction, resulting in differences in the expression of digestive enzymes and nutrient metabolism pathways. This has yet to be explored in detail, but considering that only larval stages can be infected with TSWV (Ullman 1992) and differences in digestive protein activities have been noted between adults and larvae (Yue et al. 2023)

Second, the transcriptional responses of larval salivary glands (SGs) to TSWV infection were substantially more pronounced than those observed in adults, consistent with previous proteomic analyses (Rajarapu et al. 2022). Larvae exhibited the greatest proportion of differentially expressed genes, reflecting the recent establishment of viral infection as TSWV first colonizes the SGs during the second larval instar. This acute phase of infection was accompanied by extensive transcriptional reprogramming, including suppression of ribosome biogenesis, RNA metabolism, protein synthesis, nutrient transport, detoxification, and digestive functions, as well as activation of developmental, morphogenetic, and signaling pathways. In contrast, the comparatively muted responses observed in adult SGs likely reflect prolonged virus persistence and the establishment of a more tolerant host-virus interaction following chronic infection. Finally, similarities between the transcriptomic and proteomic datasets reinforce the conclusion that cytoskeletal remodeling is a central feature of thrips–TSWV interactions. Both studies identified perturbations of cytoskeletal and muscle-associated components, suggesting that structural reorganization of SG tissues accompanies viral infection and may facilitate viral replication, intracellular movement, or transmission. Together, these complementary transcriptomic and proteomic analyses provide a more comprehensive understanding of the molecular interactions between TSWV and its thrips vector, highlighting candidate pathways involved in host shutoff, tissue remodeling, and behavioral regulation that represent promising targets for disrupting viral acquisition, persistence, and transmission within the salivary glands.

## Author Contributions (CRediT)

**Joshua Benoit:** Conceptualization, Methodology, Visualization, Formal analysis, Investigation, Data Curation, Original Draft. **Sulley Ben-Mahmoud:** Investigation, Formal analysis, Writing - Review & Editing. **Swapna Priya Rajarapu:** Formal analysis, Writing - Review & Editing. **Christopher J. Holmes:** Formal analysis, Writing - Review & Editing. **Samuel T. Bailey:** Formal analysis, Writing - Review & Editing. **Diane Ullman:** Conceptualization, Supervision, Writing - Review & Editing. **Dorith Rotenberg:** Conceptualization, Supervision, Writing - Review & Editing.

## Data availability statement

All new RNA-seq datasets for this project are associated with the NCBI Bioproject: PRJNA1311131

