## Supplementary figures and images for "Stage-specific and tomato-spotted wilt virus infection-induced changes in the salivary gland transcriptome of western flower thrips"

### Figure S1

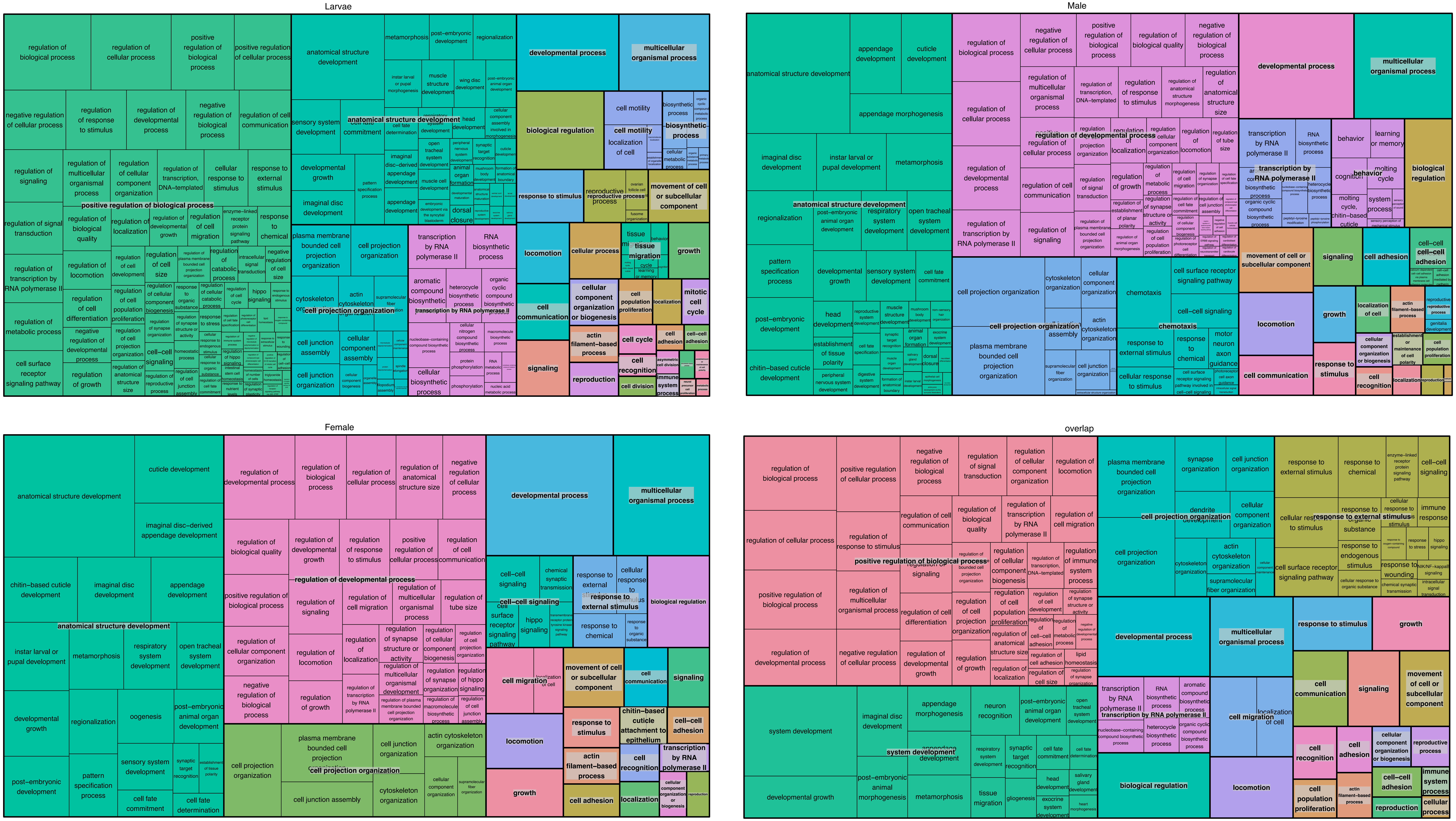

### Figure S2

#### A. Increased during TSWV infection in larvae

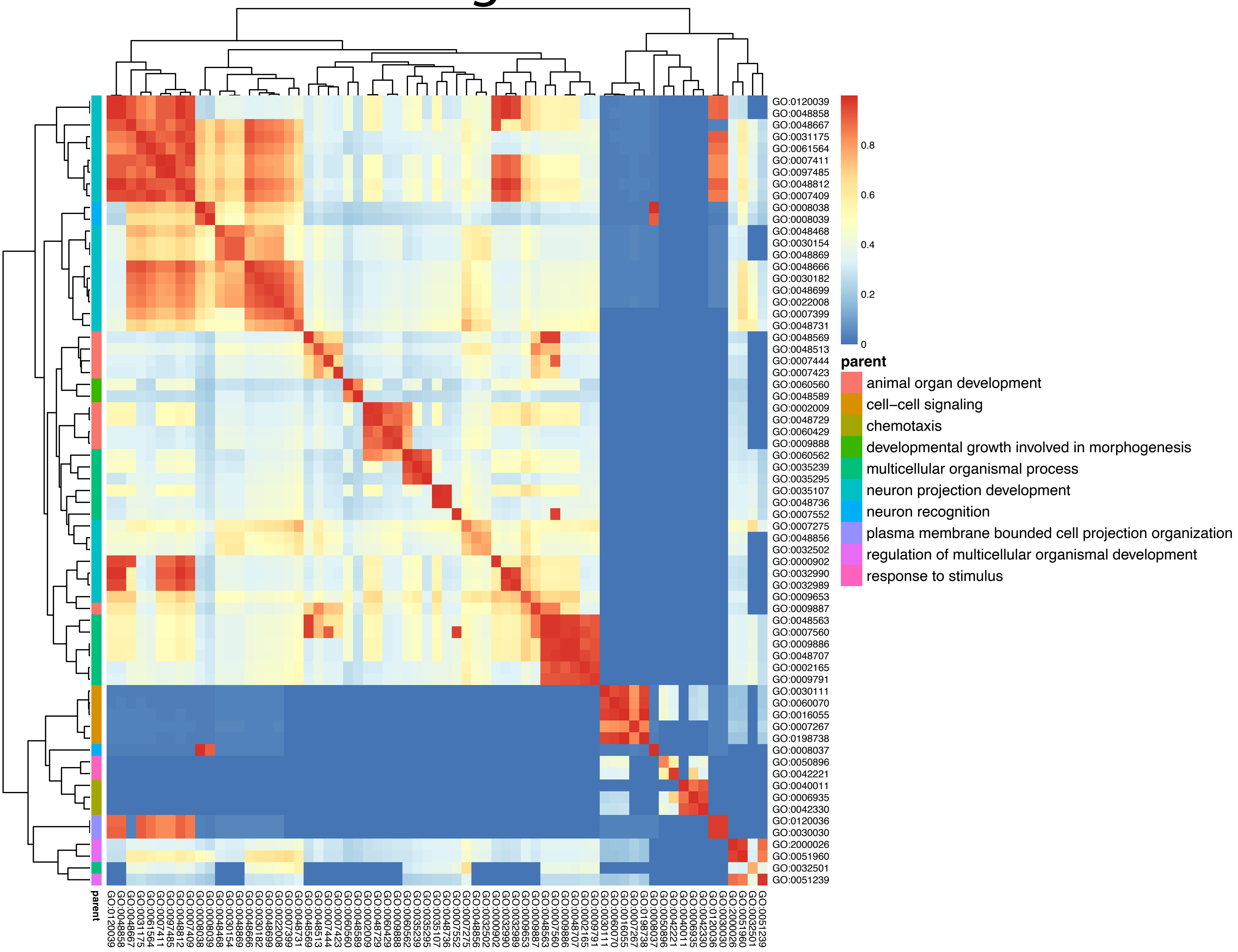


#### B. Decreased during TSWV infection in larvae

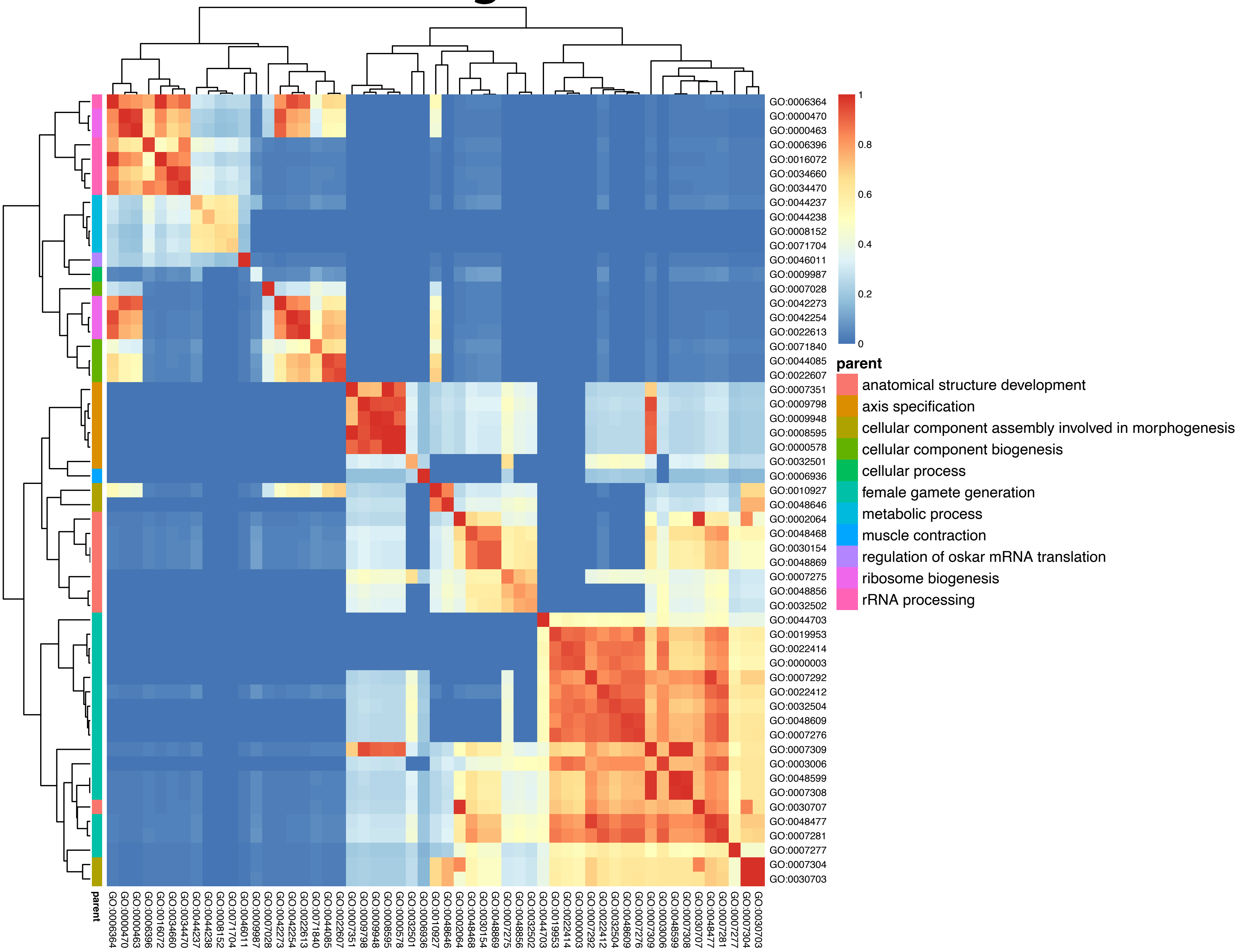
